# GPCRVS - AI-driven decision support system for GPCR virtual screening

**DOI:** 10.1101/2024.10.28.620626

**Authors:** Dorota Latek, Khushil Prajapati, Paulina Dragan, Matthew Merski, Przemysław Osial

## Abstract

**Summary:** G protein-coupled receptors (GPCRs) constitute the largest and most frequently used family of molecular drug targets. The simplicity of GPCR drug design results from their common seven-transmembrane helix topology and well-understood signaling pathways. GPCRs are extremely sensitive to slight changes in the chemical structure of compounds, which allows for the reliable design of highly selective and specific drugs. Only recently has the number of GPCR structures, both in their active and inactive conformations together with their active ligands, become sufficient to comprehensively apply machine learning in decision support systems to predict compound activity for drug design. Here, we describe GPCRVS, an efficient machine learning web service for the online assessment of compound activity against several GPCR targets, including peptide and protein-binding GPCRs, the most difficult for virtual screening tasks. As a decision support system, GPCRVS evaluates compounds in terms of their activity range, the pharmacological effect they exert on the receptor, and the binding modes they could possibly demonstrate for certain types of GPCR receptors. GPCRVS can be applied to compounds ranging from small molecules to short peptides uploaded as common chemical format files. The activity class assignment and the binding affinity prediction are provided in a reference to similar predictions precomputed for already known active ligands of each GPCR target. This multiclass classification in GPCRVS, handling often incomplete and fuzzy biological data, was validated on ChEMBL-retrieved data sets for secretin-like and chemokine GPCR receptors.

**Availability and Implementation:** GPCRVS is freely available at: https://gpcrvs.chem.uw.edu.pl

## Introduction

The majority of the human proteome is still considered undruggable or too complex to propose efficient and selective and/or specific enough therapeutics^1^. Among successful drug targets, the largest protein population belongs to ion channels, followed by G protein-coupled receptors (GPCRs), kinases, and finally nuclear receptors^1^. However, known small-molecule drugs of high bioavailability mostly target GPCRs (33%), followed by ion channels (18%), nuclear receptors (16%), and kinases (3%). The structural similarity of GPCRs and their common mode of activation during signal transduction imposes problems in drug discovery regarding unwanted off-target effects and a lack of drug selectivity. This requires the massive testing of drug candidates against many possible protein targets prior to clinical studies, leading to a highly increased drug development cost. Recent advances in artificial intelligence (AI) techniques demonstrated unexpected possibilities of accelerating the validation of drug candidates through the automatic detection of unwanted similarities of their chemical structures and modes of action. Mimicking human expertise in finding patterns responsible for the compound activity against a certain molecular target is a major advantage of AI systems currently being developed for drug design. Although such systems are often considered ‘black box’ techniques, uncovering their mechanisms of self-learning by explainable AI^2^ provides another way of looking at biological problems that is inaccessible to human experts.

Here, we describe GPCRVS – a recently developed web service for virtual screening against drug targets from class A and B GPCRs that are activated endogenously by peptides or small proteins (Figure 1). GPCRVS overcomes the limitations of component methods for virtual screening such as ligand-based or target-based approaches. In addition, GPCRVS allows for the testing of drug candidates in terms of their types (agonists vs. antagonists vs. modulators) and possible off-target effects against several well-known GPCR drug targets, including secretin-like and chemokine GPCR receptors. In particular, we propose a novel approach, 6-residue peptide truncation, to combine an overwhelming number of active peptide compounds available in ChEMBL for GPCRs together with small-molecule compounds into one complete training data set. GPCRVS was tested and compared with a number of other similar, currently accessible methods^3–5^ that already implement ML or AI systems in general for drug design.

**Figure 1.**
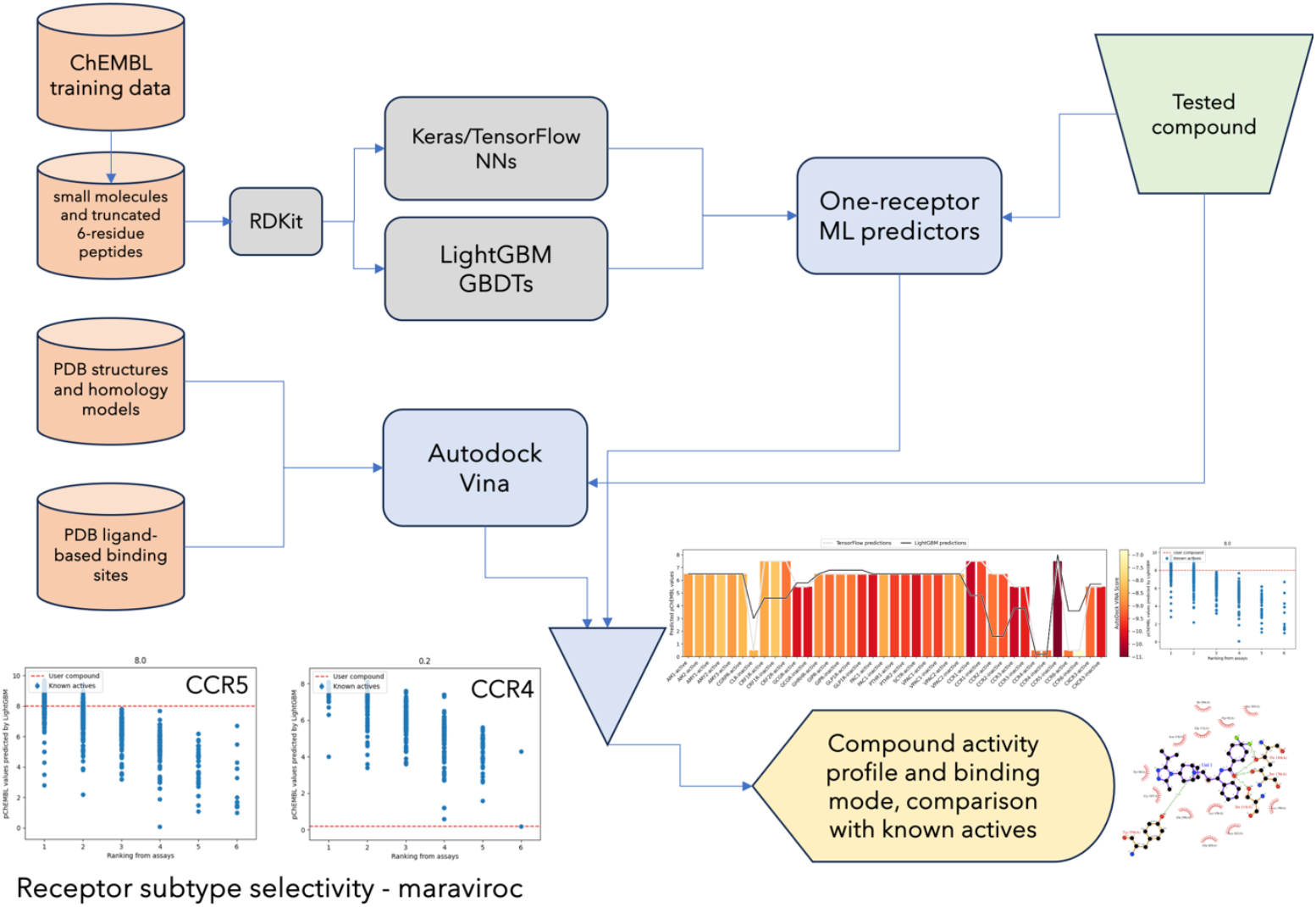
The scheme of GPCRVS describing used training data sets, implemented algorithms, and provided results. Bottom left – example results for maraviroc showing GPCRVS performance in terms of receptor subtype selectivity prediction.

## Material and methods

### Deep neural networks implemented in GPCRVS

The ligand-based virtual screening module in GPCRVS includes two diverse ML algorithms (Figure 1), deep learning neural networks implemented in TensorFlow^6,7^ and gradient boosting decision trees implemented in LightGBM^8^. Keras API was used to build deep neural networks (DNNs)^9,10^ followed by hyperparameter tuning^11^ and model evaluation in 10-fold cross-validation. The best DNN model for each receptor was implemented in GPCRVS for multiclass classification tasks (ranges of pChEMBL values^12^).

### Gradient boosting machines implemented in GPCRVS

LightGBM outperforms other gradient boosting methods due to gradient-based one-side sampling and exclusive feature bundling. It was used to build a gradient boosting machine (GBM) model with decision trees as base-learners and the RandomizedSearch optimization of hyperparameters with 10-fold cross validation that is comparable to the hyperparameter tuning by Optuna^13^. The best GBM model for each receptor was implemented in GPCRVS for compound activity prediction (pChEMBL values).

### Ligand binding mode prediction in GPCRVS

AutoDock Vina^14^, previously extensively tested against several class B GPCRs^15^ in comparison with the Schrödinger-licensed Glide^15,16^ and successfully used in^17^, was implemented in GPCRVS for molecular docking (Figure 1). Orthosteric and allosteric binding sites of agonists, antagonists and positive and negative allosteric modulators were considered separately for each receptor type (Figure 2). Receptor structures were either based on PDB structures with Maestro-based missing atom filling and refinement^18^ or, if not present, prepared with the previously described Modeller/Rosetta CCD loop modeling approach to GPCR structure prediction^17^ implemented in GPCRM^19,20^.

**Figure 2.**
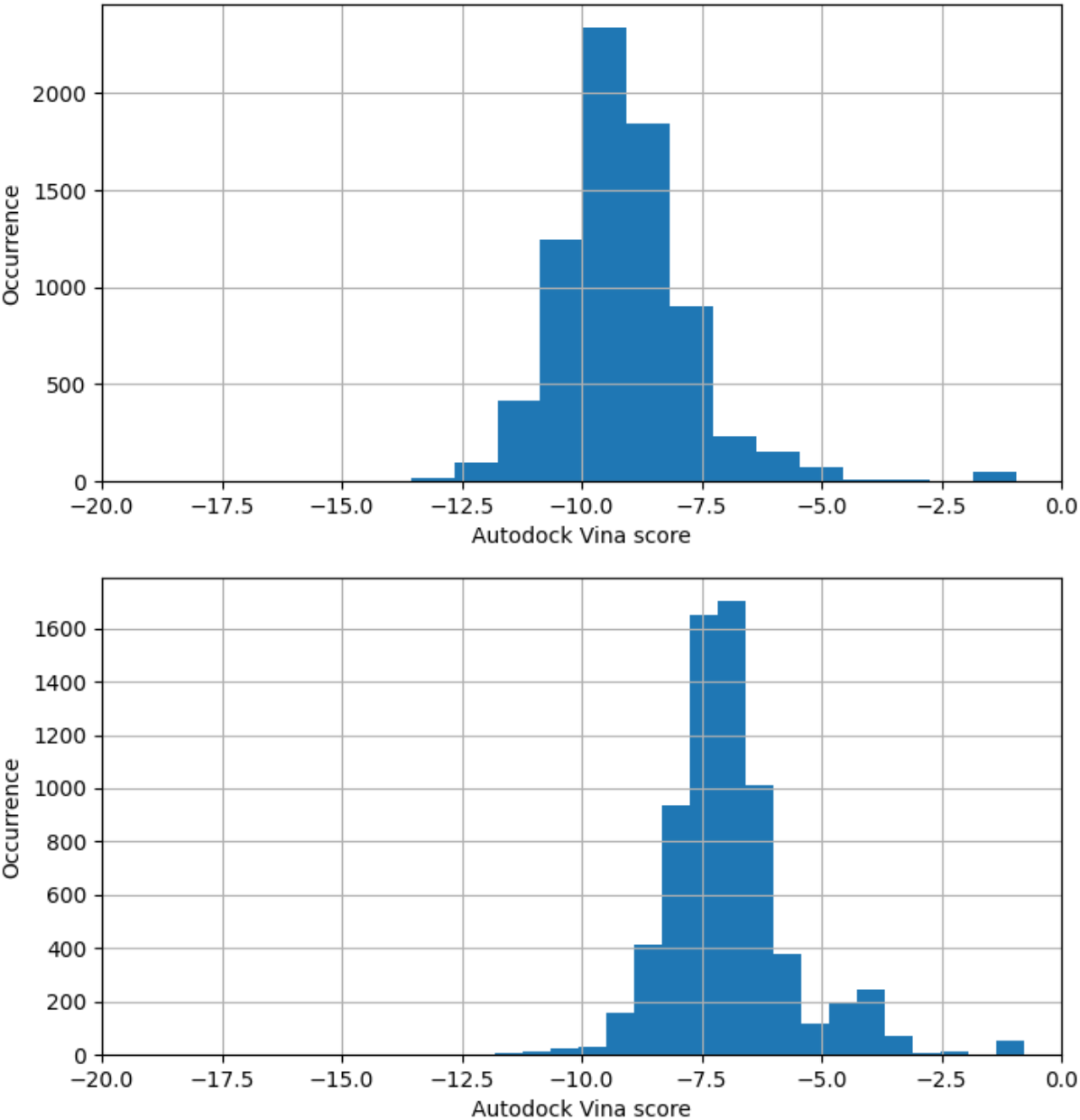
Histograms obtained for active GPCR ligands retrieved from ChEMBL showing their high affinity for their GPCR receptors as observed in molecular docking by AutoDock Vina. Top – orthosteric ligands, bottom – allosteric ligands.

### Data sets used for training and validation

Training data sets were retrieved from ChEMBL and curated in terms of uniqueness, SMILES correctness, and assay type compatibility. An 80/20% ratio between training and validation sets was applied with random splitting. Peptide compounds were truncated to activating 6-residue long N-terminal fragments, and converted to SMILES. Peptide compounds activating opioid, chemokine, and class B receptors tend to bind with their N-terminal short fragments (or close to the N-terminus^21^), which carry the activation ‘message’^22–24^, while C-terminal regions (‘addresses’) are responsible for the correct orientation, binding stability, and localizing interactions with the receptor and the ECD domain and/or lipid bilayer preceding the receptor binding^25^. Certain other receptors, such as apelin, endothelin or neurotensin receptors, have C-terminal activating fragments instead^26,27^. To retrieve compound-unique chemical features, extended connectivity fingerprints with a bond diameter equal to 4 (ECFP4) based on Morgan fingerprints were used^28,29^.

## Results

### GPCRVS as a decision support system for newly designed compounds

The typical output provided by GPCRVS includes an activity profile of a compound representing its predicted activity against several GPCR drug targets (Figure 1). There are three, uncorrelated in terms of their results (Figure 3), components of this decision support system – a DNN-based classification of compound activity to one of six activity classes, a GBM-based prediction of compound activity, and an AutoDock Vina-based prediction of the binding affinity and binding mode. The former two components rely on the pChEMBL standardization of compound activity markers such as EC50. The compound binding affinity prediction is based on a scoring function implemented in AutoDock Vina, including only van der Waals-like potential, hydrogen bonding and hydrophobic terms, and conformational entropy penalty, in contrast to much less computationally efficient, electrostatics-involving, the AMBER-based force field of AutoDock4.2^14^. In addition to their activity and off-target effect prediction, compounds are positioned in every orthosteric/allosteric binding site of all of the receptors with advanced AutoDock Vina sampling of the ligand-protein conformational space using a Monte Carlo search algorithm that includes an improved implementation of macrocycle flexibility^14^. The performance of AutoDock Vina in binding mode prediction was previously extensively tested^30^, also in case of GPCR ligands^31,32^, and proved to be slightly better in the prediction of binding poses than AutoDock4^30^.

**Figure 3.**
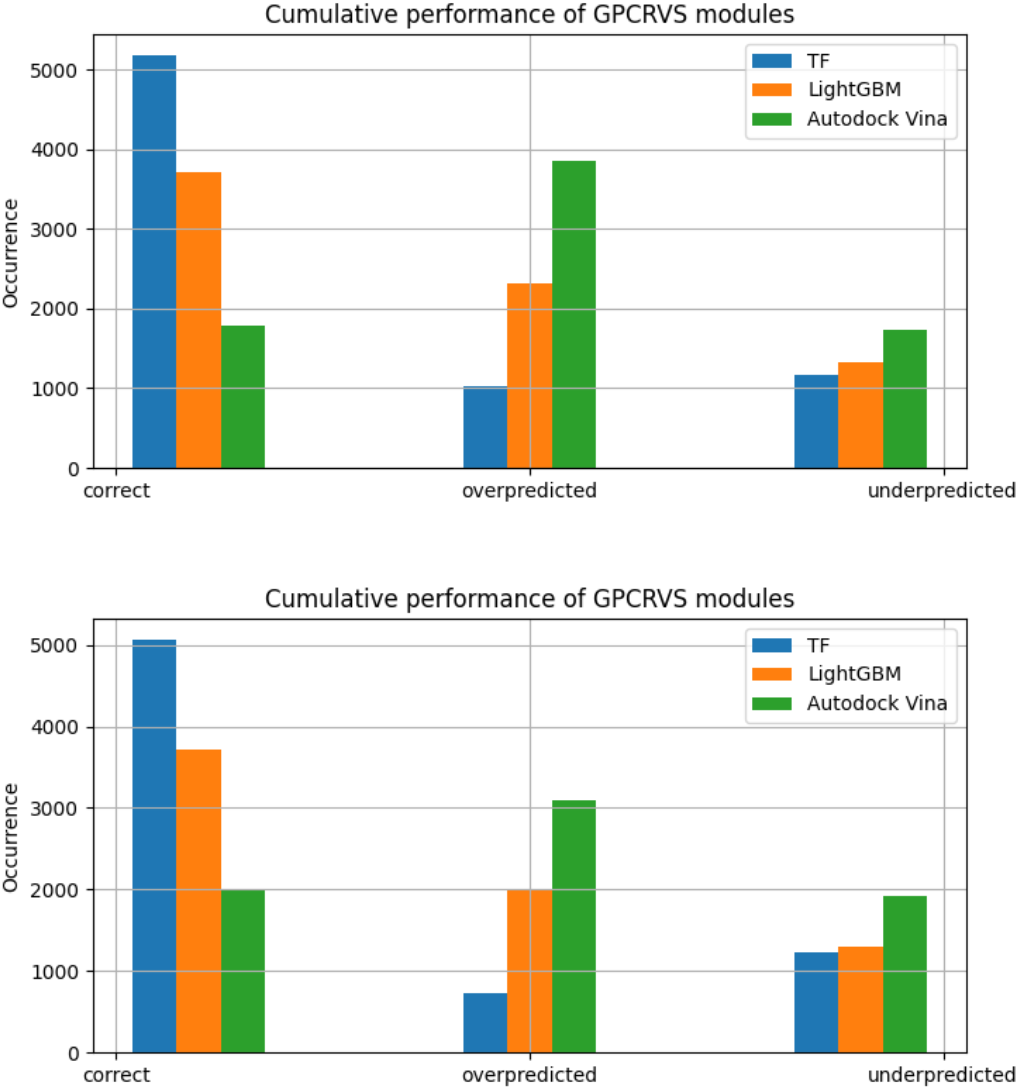
The cumulative performance of the GPCRVS modules – results for ChEMBL GPCR orthosteric (top) and allosteric ligands (bottom).

These three components of the GPCRVS decision support system represent completely different approaches to virtual screening (ligand-based vs. target-based, DNN-based vs. GBM-based, multiclass classification vs. regression). Thus, the compound activity prediction is not biased by an overlap between component methods but is solely due to the compound structure and resulting properties. Beta-version of GPCRVS achieved accuracy as high as 71.4% in the drug target prediction task if at least one component predictor was accurate and 33.3% if the condition of agreement of at least two predictors was applied (28.6% if they were only ML predictors). Although the resulting average prediction accuracy for this limited validation data set of selective GPCR ligands seems low, it is still higher than the prediction accuracy of each of the methods separately (38.1%, 47.6%, 19.0% for the DNNs, GBMs, and AutoDock Vina, respectively) and thus a significant improvement is observed compared to our previous AutoDock Vina-based web service^16^ or similar methods^33,34^. What is more, DNNs and GBMs perform uncorrelated drug target class prediction as shown by the accuracy of their individual predictions comparing the common prediction (28.6 %).

GPCRVS demonstrated high prediction accuracy for the validation data sets, added each time for comparison to predictions for a tested compound. For most of the included GPCRs, the compound activity ranking based on data from functional assays retrieved from ChEMBL correlated with the DNNs/GBMs/AutoDock Vina predictions of pChEMBL values (Figure 4). GPCRVS also performed well in the case of binary classification (active vs. inactive, Figure 5) as tested in detail by us in^12^.

**Figure 4.**
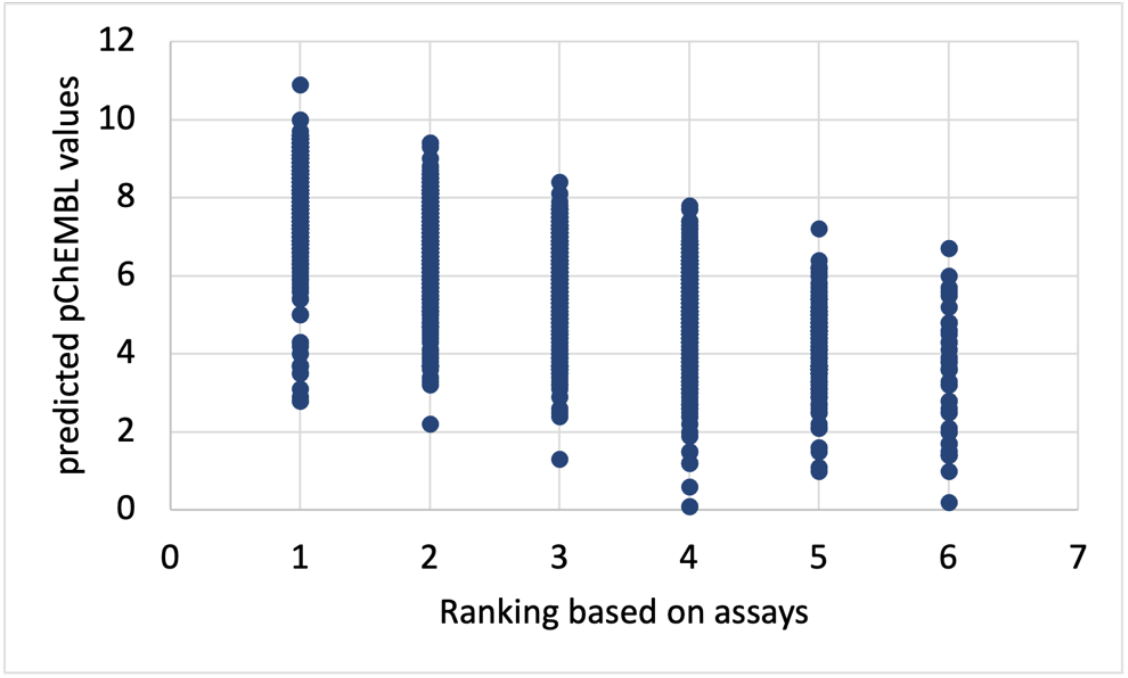
A comparison of pChEMBL values predicted by DNNs with the compound activity ranking based on results from pharmacological assays obtained from ChEMBL (the higher pChEMBL value, the higher rank assigned). DNN-based predictions correlate with the experimentally determined compound activity.

**Figure 5.**
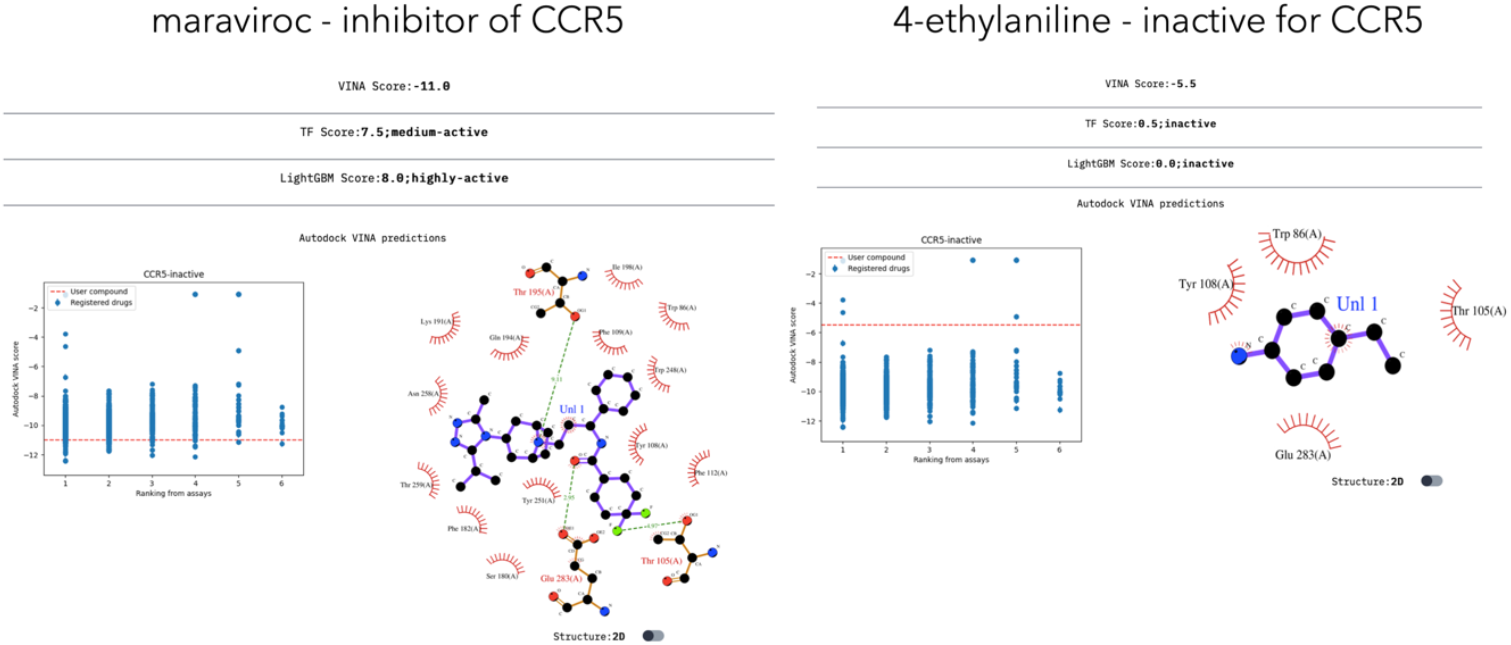
Example results of binary classification in GPCRVS - two diverse compounds (CCR5 inhibitor and an inactive compound) tested against CCR5. All three component methods predicted activity correctly.

## Conclusions

GPCRVS represents an efficient, simple, easily accessible decision support system for drug design, that aims to facilitate preclinical drug design for G protein-coupled receptors binding peptide and small proteins. In addition, a novel approach to use peptide ligands in conjunction with small-molecule ligands for the training of DNN and GBM models was proposed. This makes it possible to benefit from all GPCR ligand data sets deposited in ChEMBL, and to design new drugs that could include both peptide and non-peptide scaffolds of increased unified activity and selectivity. Currently, both rhodopsin and secretin-like, peptide/small protein-binding GPCR receptors are included in GPCRVS, allowing it to make comparative predictions between GPCR subfamilies at the same time. The added value of GPCRVS is providing three, diverse in principle, algorithms to assess compound affinity and activity for GPCR targets which indeed provide internally uncorrelated results.

## Acknowledgements

This research was funded by the National Science Centre in Poland, grant number 2020/39/B/NZ2/00584. Computational resources and maintenance are provided by Faculty of Chemistry, University of Warsaw. All packages are licensed under the Apache License, Version 2.0 (http://www.apache.org/licenses/LICENSE-2.0). Computational resources were provided by High-performance Infrastructure PLGrid (Poland), grant number PLG/2023/016255.

## Author contributions

Conceptualization, supervision, project administration, funding acquisition, resources, writing – original draft – DL; writing – review & editing – DL, PD; investigation – DL, KP, PD; methodology – DL; software – DL, KP, PO, PD; data curation – MM, DL, PD; validation – DL, PD; formal analysis – DL; visualization – DL, PO, PD.

